# Chromosome-level reference genome assembly for the protected resource plant, *Zenia insignis*

**DOI:** 10.1101/2024.06.01.596946

**Authors:** Si-Yun Chen, Zhi-Yun Yang, Rong Zhang, Hui Liu, Ting-Shuang Yi

## Abstract

*Zenia insignis* Chun, from the subfamily Dialioideae of Fabaceae, is a tree species of significant economic and ecological values. It is a threatened species as per the IUCN Red List. The absence of a reference genome has impeded research on *Z. insignis* despite its importance. In this study, we generated a reference genome for *Z. insig n*, *i*w*s* ith a contig N50 of 6.02Mb, a total length of 361.11 Mb, and 97.71% of the sequences assembled into 14 pseudo-chromosomes. The BUSCO assessment score for completeness is 97.30%, and the LAI index assessment score for continuity is 14.57. To our knowledge, this is the first reported nuclear genome of a species belonging to subfamily Dialioideae. The reference genome will provide a valuable resource for the phylogenomic studies of the Fabaceae family and facilitate further research on *Z. insignis*.

## Background & Summary

*Zenia insignis* Chun, a member of the family Fabaceae, is a tree species recognized as one of the Grade-II Wild Plants of National Priority Protection in China^1^ and is listed as Near Threatened (NT) species on the International Union for Conservation of Nature (IUCN) Red List^2^. This species was named by the renowned botanist Woon-Young Chun in honor of Hongjun Ren (H.C. Zen)^3^. The species is primarily distributed in southern and southwestern China at an altitude range of 200–1000 meters and northern Vietnam^4^. *Zenia insignis* boasts a robust root system and strongly resists drought and infertile conditions^5^. In the karst regions of southern and southwestern China, it has been utilized as an outstanding tree species for afforestation on rocky terrain and as a pioneering species for the control of rocky desertification^5–7^. Additionally, it has significant economic value. The wood of *Z. insignis* is versatile and suitable for producing doors, boards, agricultural implements, and furniture^8,9^. The plant’s tender leaves serve as livestock feed and green manure in rice fields^7–9^.

Research on *Z. insignis* has predominantly concentrated on seedling^10,11^, reforestation technologies^5,6^, and phylogenetic studies^12–19^. Though partial and complete plastome sequences^12–14,16^ and nuclear sequences^15,17–19^ have been utilized in constructing the phylogenetic tree of the Fabaceae family, the deep phylogenetic relationships within Fabaceae remain to be fully elucidated. A chromosome-level genome sequence of *Z. insignis* would be instrumental in clarifying these phylogenetic relationships. In this study, we have successfully obtained the reference genome of *Z. insignis* and performed comparative analyses with other representative plant genomes to further our understanding of the evolutionary history of the Fabaceae.

In this study, we integrated multiple sequencing data types to assemble and annotate the genome of *Z. insignis*. Specifically, we used PacBio continuous long reads (CLR) (174.14 Gb, 363.43× coverage), next-generation sequencing (NGS) reads (44.14 Gb, 91.96× coverage), Hi-C data (59.63 Gb, 124.45× coverage), and full-length transcriptome reads. The resulting reference genome consists of 14 pseudo-chromosomes and spans a total length of 352.84 Mb with a contig N50 of 6.02 Mb. The genome’s LAI index score is 14.57. The GC content of the genome is 34.38%, and the proportion of repeat sequences is 35.47%, with long terminal repeat (LTR) elements being the most prevalent, accounting for 9.03%. We also obtained the chloroplast genome sequence of *Z. insignis* with a length of 159,390 bp.

We predicted 33,322 gene models, of which 90.46% (30,143) were functionally annotated. Analysis of orthologous genes between *Z. insignis* and ten other representative plant species revealed 608 orthogroups, with at least 72.72% of species having single-copy genes in any given orthogroup. Additionally, 1,625 gene families underwent significant expansion, and 741 gene families underwent significant contraction in *Z. insignis*. In the whole genome duplication analysis, two collinear block *Ks* peaks were observed at 0.245±0.001 and 1.693±0.001, suggesting that *Z. insignis* has undergone an additional whole genome duplication (WGD) event after the γ WGT event shared by all core eudicots.

## Methods

### Genome size estimation and sequencing

Fresh young leaves of *Z. insignis* were collected from the Kunming Botanical Garden (KBG), Kunming Institute of Botany, Chinese Academy of Sciences (KIB), Yunnan Province, China. The voucher specimen (Collection Number: Yi20005) had been deposited at the Herbaria (KUN) of Kunming Institute of Botany, Chinese Academy of Sciences, Yunnan Province, China. The leaves were immediately flash-frozen in liquid nitrogen to preserve their cellular structure and molecular content. Subsequently, the samples were transferred to an ultra-low temperature freezer maintained at –80°C for storage.

DNA/RNA extraction, library construction, and sequencing processes were provided by Wuhan Frasergen Bioinformatics Co., Ltd. The major steps are summarized below:

1. Genomic DNA was extracted from the tender leaves of *Z. insignis* using the modified cetyltrimethylammonium bromide (CTAB) method^20,21^. Total RNA was isolated using a Trizol reagent (Invitrogen, CA, USA). The quantity and quality of the DNA/RNA were determined using a NanoDrop 2000 spectrophotometer (NanoDrop Technologies, Wilmington, DE, USA).
2. The quality-compliant DNA was further tested for purity and integrity using 0.8% agarose gel electrophoresis and a Qubit dsDNA HS Assay Kit on a Qubit 3.0 Fluorometer (Life Technologies, Carlsbad, CA, USA). RNA contamination was identified using a 1.5% agarose gel, and purity and integrity were evaluated using a NanoDrop 2000 and a Bioanalyzer 2100 system (Agilent Technologies, CA, USA).
3. According to the manufacturer’s instructions, a short-read library was constructed using the VAHTS Universal DNA Library Prep Kit (Vazyme, Nanjing, China) for MGI. The library’s quantification and sizing were measured using a Qubit 3.0 Fluorometer and a Bioanalyzer 2100 system (Agilent Technologies, CA, USA). Paired-end sequencing of the short-read library was conducted on an MGI-SEQ 2000 platform (BGI, China).
4. To perform long-read sequencing, the library was prepared in accordance with the PacBio single-molecule real-time (SMRT) protocol and then sequenced on a Pacific Biosciences Sequel II System in CLR mode.
5. The construction of the Hi-C library commenced with the cross-linking of fresh leaves of *Z. insignis* under vacuum infiltration with 3% formaldehyde. Following lysis, digestion, labelling, ligation, purification, adaptor addition, and PCR amplification, the Hi-C libraries were quantified and sequenced on the MGI-SEQ 2000 platform.
6. For short-read transcriptome sequencing, RNA was isolated from Z. insignis young leaves using the QIAGEN kit according to the manufacturer’s instructions. The short-read library was subsequently constructed and sequenced on the MGI-SEQ 2000 platform using paired-end sequencing.
7. The full-length transcriptome sequencing was conducted using the Pacific Biosciences DNA Template Prep Kit 2.0 for library construction (Pacific Biosciences, CA, USA). Subsequently, the SMRT sequencing was performed on a Pacific Biosciences Sequel II platform.

Our study generated 44.14 Gb of raw data for the whole genome, 174.14 Gb of PacBio CLR data, 59.63 Gb of Hi-C data, and 81.92 Mb of full-length transcriptome data (Table 1).

**Table 1.** Statistics of sequencing data.

| Survey data | PacBio CCS data | Hi-C data | Full-length<br>Transcriptome data |
| --- | --- | --- | --- |
| 44140.85Mb | 174,139.06Mb | 59,630.13Mb | 81.92Mb |
| 91.96× | 363.43× | 124.45× | — |

The genome size of *Z. insignis* was assessed using flow cytometry analysis conducted on a BD FACScalibur flow cytometer (BD Biosciences). The test sample preparation was conducted in accordance with the manufacturer’s guidelines and aligned with established methodologies from previous literature^22,23^. As a standard reference, we utilized *Zea mays* (maize)^24^, a species with a well-documented genome size of 2.3 Gb. The analysis yielded a result indicating that the genome size of *Z. insignis* is approximately 480 ± 4.8 Mb (Figure 1A).

**Figure 1.**
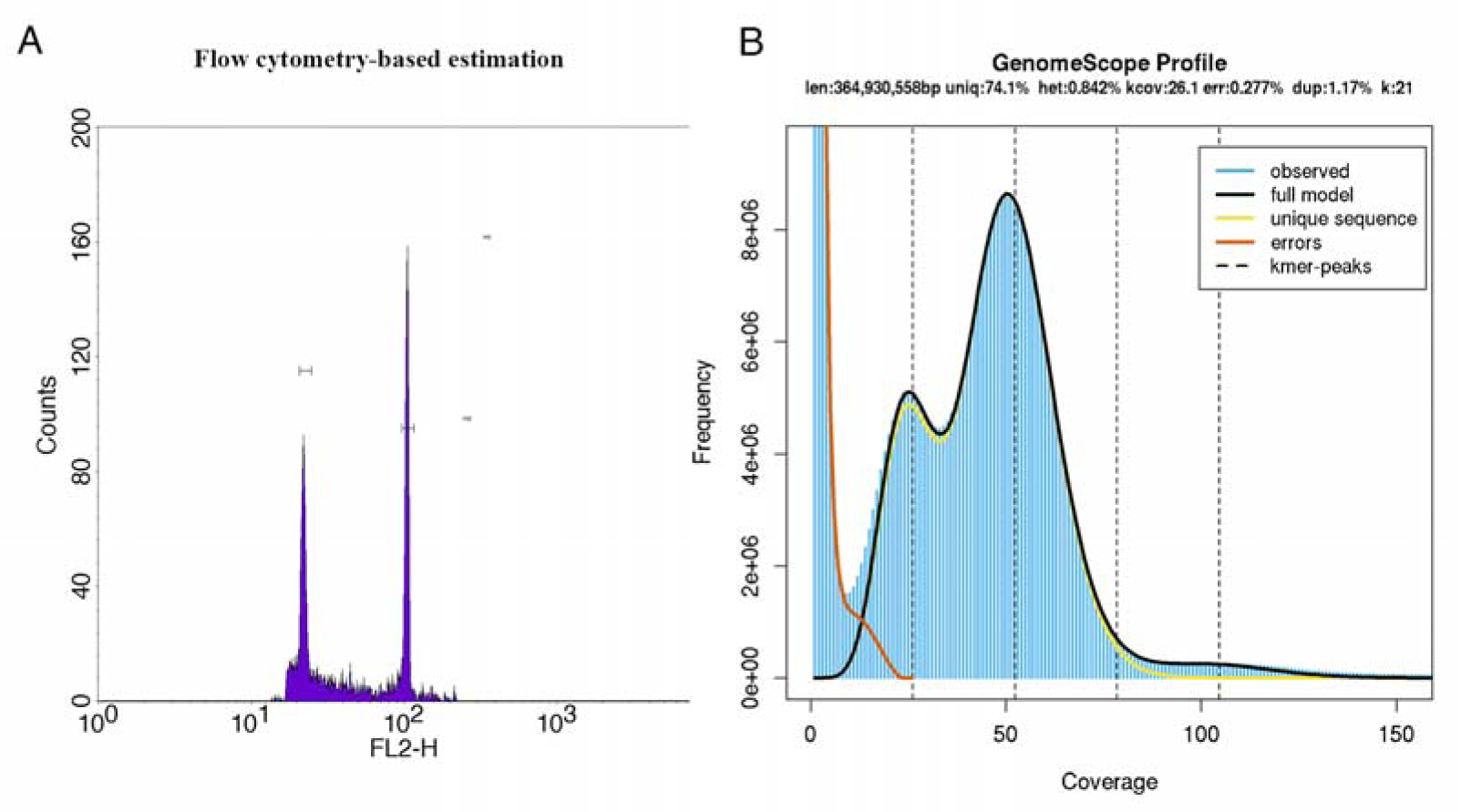
The result of genome size estimation for *Z. insignis*. (A) Flow cytometry-based estimation result of *Z. insignis* (left peak, the sample) and *Zea mays* (right peak, the reference). (B) The 21-*kmer* distribution estimation result was obtained using GenomeScope2. The distribution of *21-kmer* exhibited twoh major peaks, indicating certain heterozygosity within the genome, and high-frequency *k-mers* with coverage near 1 that was possibly generated by sequencing errors.

A *K-mer* analysis was conducted on *Z. insignis* to estimate its genome size. Next-generation sequencing (NGS) data was obtained from Frasergen and processed Fastp v0.19.3^25^. This involved the removal of adapter sequences, the filtration of reads deemed to be of insufficient length, and the elimination of those with low quality. Subsequently, Jellyfish v2.2.10^26^ was employed to calculate the frequency distribution of the cleaned data using 21-*mers*. Subsequently, GenomeScope v2.0^27^ was employed to estimate the fundamental characteristics of the genome. The genome size of *Z. insignis* was estimated to be approximately 364.94 Mb, with a heterozygosity rate of 0.842% and a repeat content of 1.17% (Figure 1B).

### *De novo* genome assembly

Figure 2 illustrates the methodology employed for the chromosome-level genome assembly of *Z. insignis*. The objective was to utilize three distinct assemblers, compare their resulting assemblies, and achieve a high-quality assembly. The assemblers employed were Falcon v0.3.0^28^, Canu v2.1.1^29^, and Flye v2.9^30^. The following section provides a summary of the assembly process.

1. The Falcon software was employed to assemble the CLR long reads. The resulting contigs were then polished using long reads with GCpp v2.0.0^31^. Subsequently, the processes above yielded the assembly contig, designated as Assembly_Contig1. Subsequently, Assembly_Contig1 was subjected to further refinement with short reads, employed NextPolish v1.31^32^. The output from the process was designated as the ‘Falcon contig’. To eliminate redundancy, the Falcon contig was processed with Purge_Dups v1.2.5^33^, and the resulting non-redundant contigs were designated as the ‘Falcon_purge contigs’.
2. The Canu software was employed to assemble the CLR long reads, and the result was subsequently polished with long reads by GCpp. Through these processes, we obtained Assembly_Contig2. Assembly_Contig2 was then polished with short reads using NextPolish, with the output from this step designated as the ‘Canu_contig’. The Canu_contig was subsequently subjected to further processing to eliminate redundancy, with the result that the non-redundant contigs were designated as the ‘Canu_purged contigs’. Purge_Dups also employed the process of eliminating redundancy.
3. The third assembler employed was Flye, which accepted the error-corrected reads generated by Canu as input. The output from Flye was polished with long reads by GCpp, resulting in the generation of Assembly_Contig3. Subsequently, the contig was polished with short reads using NextPolish, resulting in the ‘Flye_contig’. The Flye_contig was subjected to a process of redundancy elimination using Purge_Dups, resulting in the generation of a set of non-redundant contigs designated as the ‘Flye_purge contigs’.
4. Six distinct results were obtained after completing the three steps above. The following contigs were identified: Falcon contig, Canu contig, Flye_contig, Falcon_purge contig, Canu_purge contig, and Flye_purge contig.

**Figure 2.**
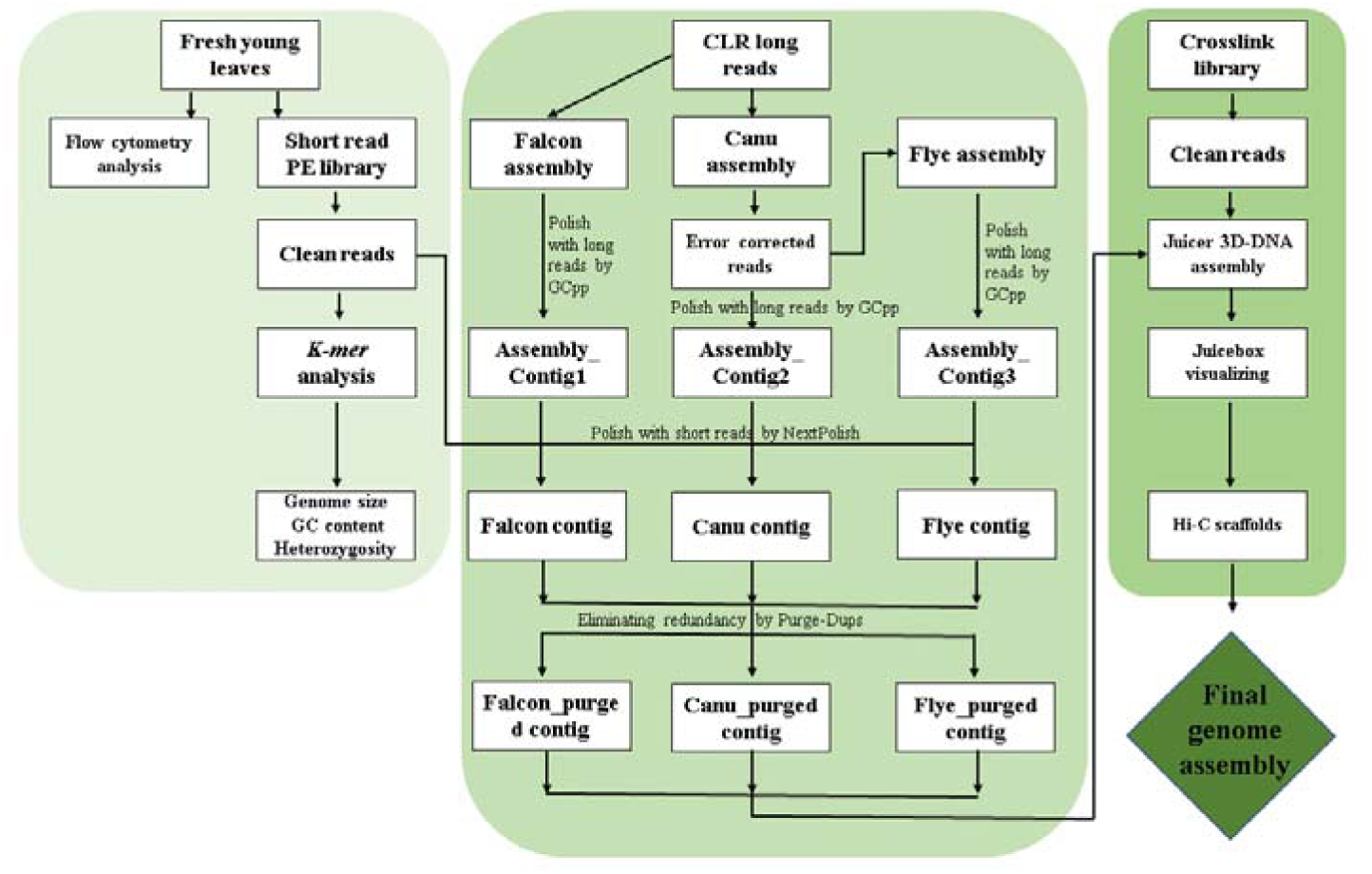
The pipeline for the chromosome-level genome assembly of *Z. insignis*.

BUSCO v5.4.5^34^ with 1614 embryophyta single-copy orthologs was employed to assess the completeness of the assembly, the N50^35^ metric provided insights into the distributions of contigs, and the LTR Assembly Index (LAI) v2.9.0^36^ was employed to evaluate the contiguity of the assembled genome. The results are presented in Tables 2–4. Notably, the Canu_purged assembly exhibited the highest N50 value, the Flye_purged assembly achieved the optimal BUSCO score, and the Falcon_purged contig assembly exhibited the highest LAI score. In other words, the result generated by Purge_Dups exhibited a better contiguity. Manual inspection using Hi-C reads revealed a low redundancy in Falcon_purge assembly (Supplementary Figure 1). Consequently, the results generated by Purge_Dups were selected for further comparison. A comparison of the Falcon_purge assembly with other results generated by Purge_Dups revealed that it exhibited high contiguity with N50 of 6.02 Mb, BUSCO completeness of 97.30%, and LAI score of 14.57.

**Table 2.** The results of the assessment for N50.

| Nx | Canu_contig |  | Canu_purged_contig |  | Falcon_contig |  | Falcon_purged_contig |  | Flye_contig |  | Flye_purged_contig |  |
| --- | --- | --- | --- | --- | --- | --- | --- | --- | --- | --- | --- | --- |
|  | Size | Number | Size | Number | Size | Number | Size | Number | Size | Number | Size | Number |
| N90 | 92,660 | 1,400 | 1,431,948 | 43 | 363,551 | 152 | 1,503,027 | 68 | 172,335 | 253 | 309,802 | 192 |
| N80 | 113,319 | 822 | 4,428,622 | 31 | 1,150,475 | 87 | 2,685,811 | 49 | 529,601 | 137 | 653,152 | 114 |
| N70 | 156,297 | 367 | 7,093,928 | 23 | 2,280,795 | 59 | 3,819,453 | 38 | 952,759 | 84 | 1,139,158 | 72 |
| N60 | 320,232 | 71 | 8,669,316 | 18 | 3,626,040 | 43 | 4,916,748 | 29 | 1,478,745 | 53 | 1,600,360 | 46 |
| N50 | 4,693,084 | 29 | 10,026,892 | 14 | 4,819,753 | 31 | 6,024,120 | 22 | 2,345,742 | 32 | 2,516,555 | 29 |
| N40 | 7,292,602 | 20 | 12,366,826 | 11 | 5,750,257 | 23 | 7,011,975 | 17 | 3,977,660 | 20 | 4,128,350 | 18 |
| N30 | 10,196,961 | 13 | 14,068,457 | 8 | 7,098,875 | 16 | 7,310,681 | 12 | 6,920,119 | 12 | 7,140,693 | 11 |
| N20 | 14,068,457 | 8 | 14,855,825 | 5 | 7,527,208 | 9 | 8,509,734 | 7 | 9,355,035 | 7 | 9,355,035 | 7 |
| N10 | 15,123,523 | 4 | 15,261,855 | 3 | 9,877,742 | 4 | 11,240,351 | 3 | 9,936,040 | 4 | 10,281,987 | 3 |
| TotalLength | 592,944,931 | 2,337 | 377,600,313 | 448 | 457,757,591 | 598 | 360,998,718 | 266 | 371,463,918 | 1,303 | 354,656,638 | 744 |

**Table 3.** The results of the assessment for BUSCO.

|  | Canu_contig | Canu_purged_contig | Falcon_contig | Falcon_purged_contig | Flye_contig | Flye_purged_contig |
| --- | --- | --- | --- | --- | --- | --- |
| Complete BUSCOs (C) | 98.50% | 98.50% | 97.50% | 97.30% | 98.60% | 98.60% |
| Complete and single-copy BUSCOs (S) | 44.10% | 77.30% | 65.80% | 79.80% | 76.70% | 76.90% |
| Complete and duplicated BUSCOs (D) | 54.40% | 21.20% | 31.70% | 18.40% | 21.90% | 21.70% |
| Fragmented BUSCOs (F) | 0.30% | 0.60% | 1.00% | 1.20% | 0.40% | 0.60% |
| Missing BUSCOs (M) | 1.20% | 0.90% | 1.50% | 1.50% | 1.00% | 0.80% |
| Total BUSCO groups | 1614 | 1614 | 1614 | 1614 | 1614 | 1614 |

**Table 4.** The results of the assessment for LAI.

|  | Chromosome<br>From | To | Intact | Total | raw_LAI | LAI |
| --- | --- | --- | --- | --- | --- | --- |
| Canu_contig | 1 | 592,944,931 | 0.0088 | 0.2003 | 4.38 | 0 |
| Canu_purged_contig | 1 | 377,600,313 | 0.0126 | 0.2099 | 5.98 | 11.61 |
| Falcon_contig | 1 | 457,757,591 | 0.0077 | 0.1684 | 4.57 | 4.62 |
| Falcon_purged_contig | 1 | 360,998,718 | 0.0119 | 0.1329 | 8.94 | 14.57 |
| Flye_contig | 1 | 371,463,918 | 0.0109 | 0.1868 | 5.83 | 11.46 |
| Flye_purged_contig | 1 | 354,656,638 | 0.0108 | 0.1689 | 6.4 | 12.03 |

During the chromosome assembly phase, we integrated Hi-C data with the previously assembled contigs to achieve chromosome-level scaffolding. Specifically, Juicer was employed to align the Hi-C reads to the contig genome. Subsequently, the 3D-DNA pipeline^37^ was used to map the contigs onto chromosome-level scaffolds. For a visual representation of the Hi-C data, Juicebox v1.22^37^ was employed. The results of the chromosome assembly also demonstrated that the Purge_Dups results exhibited lower redundancy, compared to the rest two assemblies (Supplementary Figure 1). The Purge_Dups result was chosen as the optimal choice, and subsequently, the heatmap of the result was manually adjusted (Figure 3A). The final outcome of this process was the generation of 14 pseudo-chromosomes with a total length of 352.84 Mb, as illustrated in Figures 3A and 3B.

**Figure 3.**
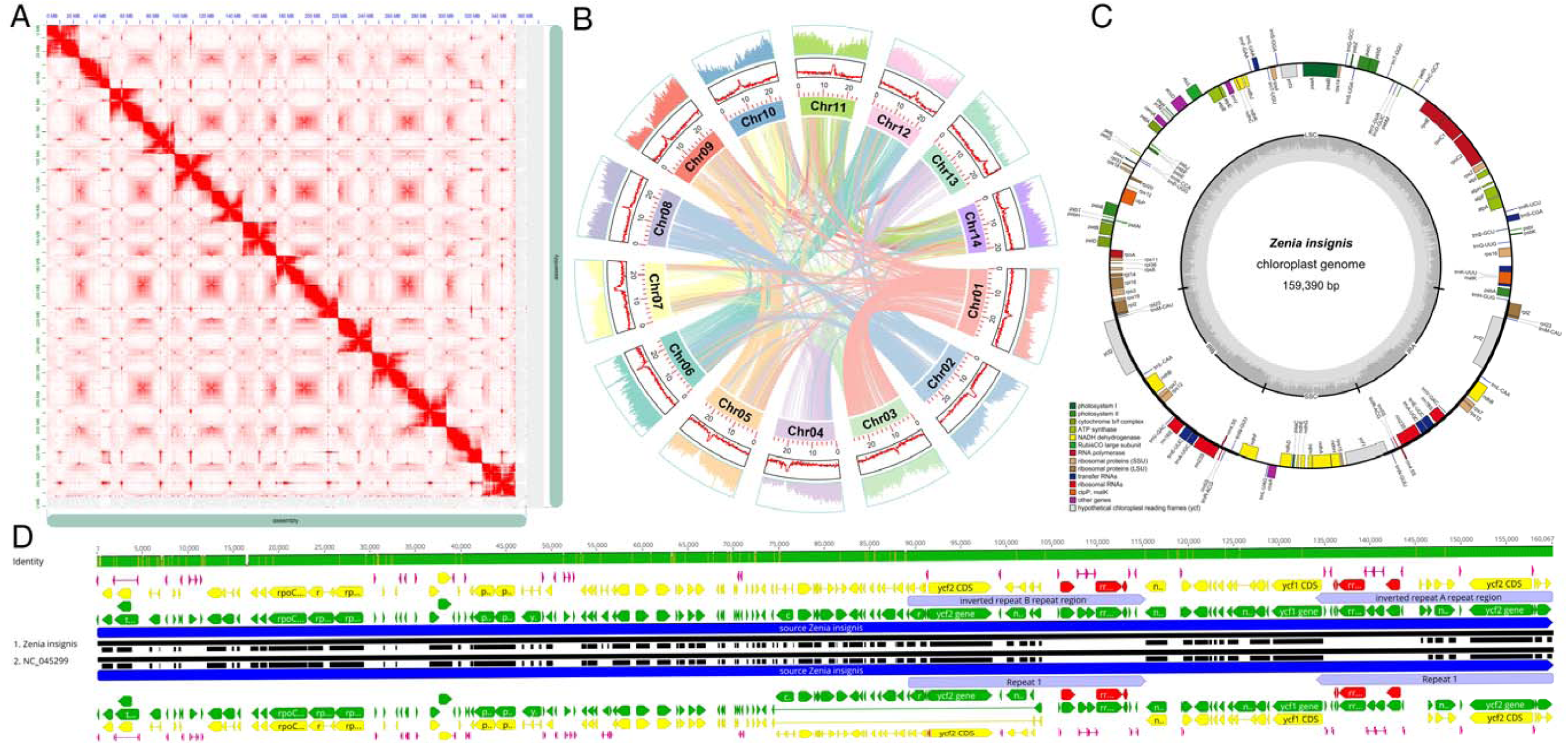
The chromosome and plastome information of *Z. insignis*. (A) The interaction heat map of Hi-C among chromosomes for the *Z. insignis* genome. The Hi-C interaction matrix shows the pairwise correlations among the 14 pseudomolecules. (B) The high-quality genomic landscape of *Z. insignis*. The genome diagram has four circles from the outside: gene density, GC content, chromosomal length, and intraspecies collinearity. (C)The visualization of the plastome genome of *Z. insignis* using OGDRAW. The outside circle displays genes color-coded according to their functional groups. The second circle illustrates the large single-copy (LSC), inverted repeat (IRA, IRB), and small single-copy (SSC) regions. The third circle delineates the percentage of GC content and genomic positions in kilobase pairs. (D) Compared analysis of plastome for *Z. insignis*. Gene(green), CDS (yellow), RNA (red), and IR regions (grey) have been color-coded according to their functional groups. The identity of the two plastome sequences exhibited high overall similarity, except for regions in non-coding sequences.

The GetOrganelle v1.7.3.3^38^ software was employed to assemble the chloroplast genome, successfully acquiring the complete chloroplast genome sequence depicted in Figure 3C. The newly assembled chloroplast genome spans 159,390 base pairs (bp) and is annotated using the PGA tool^39^, as shown in Figure 3C, drawn by OGDRAW v1.3.1^40^. For comparative analysis of the chloroplast genome, we employed the MAFFT plugin within the Geneious Prime software suite^41^ to align our sequence with a published chloroplast genome of *Z. insignis*^16^ (NC_045299.1) obtained from the National Center for Biotechnology Information(NCBI). As illustrated in Figure 3D, the alignment result revealed that apart from some single nucleotide polymorphisms (SNPs) in non-coding regions, there were no significant differences in large-scale sequences.

### Repeat annotation

The RepeatModeler v2.0.1^42^ software was employed to predict the occurrence of repeat sequences within the genome of *Z. insignis*. RepeatModeler integrates three *de novo* repeat-finding programs: RECON v1.08^43^, RepeatScout v1.0.6^44^, and LTRHarvest v5.9^45^/LTR_retriever v2.9.0^46^. For our analysis, we employed rmblast v2.13.0^47^ as the search engine. Subsequently, the output from RepeatModeler was processed using RepeatMasker v4.1.4^48^ to identify and mask the repetitive elements within the *Z. insignis* genome.

#Table 5 presents the results of the repeat sequence analysis for *Z. insignis*. A total of 128.08 Mb was identified as repetitive sequences, constituting 35.47% of the pseudo-chromosome sequences. The analysis revealed four primary classes of transposable elements within the genome: 9.03% for long terminal repeats (LTRs), 3.08% for DNA elements, 1.81% for long interspersed nuclear elements (LINEs), and 0.08% for short interspersed nuclear elements (SINEs).

**Table 5.**
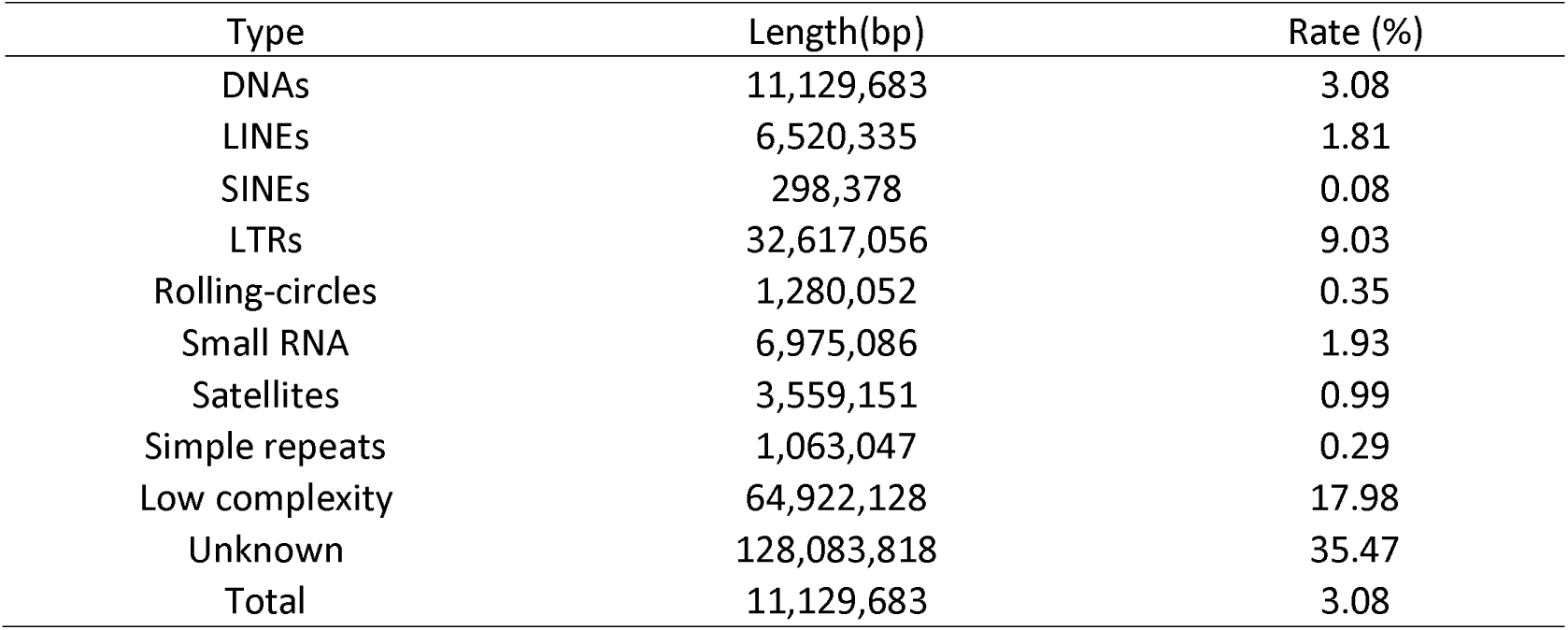
The result of repeat sequences annotation.

### Gene prediction and functional annotation

A hybrid approach was implemented to achieve high-quality gene prediction, incorporating *de novo* prediction, homology-based prediction, and RNA-Seq-assisted prediction. We employed a similar pipeline to the AnnoSmk v1.0^49^ pipeline for gene prediction. Firstly, the RepeatMasker v4.1.4^48^ software was employed to generate repeat evidence. For homology-based prediction, protein data for *Arabidopsis thaliana*^50^ was sourced from The Arabidopsis Information Resource (TAIR) and employed to construct homologous gene models using genBlastG v1.38^51^ as protein evidence. For transcriptome evidence, HISAT2 v2.2.1^52^ was employed to map paired-end short transcriptome reads to the *Z. insignis* genome. Then, the mapped reads were assembled using Trinity v2.11.0^53^. Concurrently, we processed full-length transcriptome data with the IsoSeq3 v3.4.0^54^ pipeline to produce high-quality consensus transcript sequences. Minimap2 v2.22^55^ was employed to align both short and long-read data, from which the transcriptome evidence was derived. The integrated repeat, protein, and transcript evidence were then subjected to gene prediction using MAKER2 v3.01.03^56^. Three rounds of MAKER2 analysis were executed. In the second and third rounds, the output generated through *de novo* gene prediction using AUGUSTUS v3.4.0^57^ was also included. BUSCO was employed to evaluate the gene prediction results after each round of MAKER2, yielding scores of 46.60%, 96.60%, and 96.80%, respectively (Table 6). Following the third round of MAKER2, the results were input into the PASA pipeline v2.5.0^58^, resulting in the final output. The final BUSCO score for the AnnoSmk pipeline was 97.20% (Table 6). In conclusion, a total of 33,322 gene models were predicted.

**Table 6.**
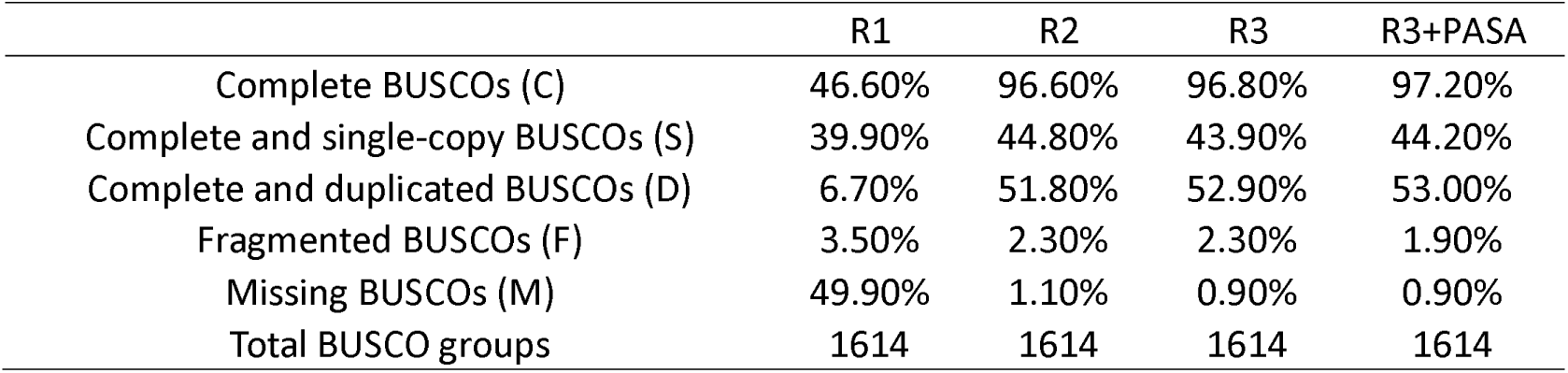
The result of BUSCO for gene prediction.

|  | R1 | R2 | R3 | R3+PASA |
| --- | --- | --- | --- | --- |
| Complete BUSCOs (C) | 46.60% | 96.60% | 96.80% | 97.20% |
| Complete and single-copy BUSCOs (S) | 39.90% | 44.80% | 43.90% | 44.20% |
| Complete and duplicated BUSCOs (D) | 6.70% | 51.80% | 52.90% | 53.00% |
| Fragmented BUSCOs (F) | 3.50% | 2.30% | 2.30% | 1.90% |
| Missing BUSCOs (M) | 49.90% | 1.10% | 0.90% | 0.90% |
| Total BUSCO groups | 1614 | 1614 | 1614 | 1614 |

The functional annotation of protein-coding genes was conducted using the eggNOG-mapper website^59^, which employs the eggNOG-mapper v2.1.12^60^ and the eggNOG 5 database. This tool utilizes several comprehensive databases, including Pfam^61^, InterPro^62^, Gene Ontology (GO)^63^, and the Kyoto Encyclopedia of Genes and Genomes (KEGG)^64^, to annotate protein sequences. This analysis resulted in the successful functional annotation of 30,143 protein-coding genes, representing 90.46% of the total gene models.

### Comparative genomics and phylogenetic analysis

The identification of orthologous genes represents a central objective in the fields of comparative genomics and phylogenetics. In this study, we compared the protein sequences of *Z. insignis* with those of ten other representative plant species, including *Arabidopsis thaliana*^50^(GCA_000001735.2), *Cercis chinensis*^65^(PRJCA007288), *Eriobotrya japonica*^66^(GWHAAZU00000000.1), *Glycine max*^67^(GCA_000004515.5), *Lysidice rhodostegia*^68^, *Medicago truncatula*^69^(GCA_003473485.2), *Senna tora*^70^(GCA_014851425.1), *Quercus mongolica*^71^(GCA_011696235.1), *Vitis vinifera*^72^(GCA_030704535.1), and *Polygala tenuifolia*^73^(TCMPG20196). These were employed to discern orthologs and paralogs. The OrthoFinder v2.5.4^74^ software was used to predict orthologous and paralogous gene families. The protein sequences from the eleven species were input into OrthoFinder, resulting in the clustering of 39,196 genes into 30,977 gene families. The analysis revealed 8,698 orthogroups that are shared by all species. Of these, 40 orthogroups comprise single-copy genes, while 608 orthogroups have at least 72.72% of species exhibiting single-copy genes within each orthogroup.

We used the 608 orthogroups obtained from our analysis to construct phylogenetic trees. Initially, the seqkit tool v0.16.1^75^ was employed to extract the corresponding nucleic acid sequences. Subsequently, the DNA sequences were aligned using MUSCLE v3.81511^76^, and the sequences were concatenated using seqkit to generate a supermatrix of genes. The supermatrix was subsequently employed as the input for constructing the phylogenetic tree, which was generated using RAxML v8.2.12^77^ with the maximum likelihood method. The most appropriate model for RAxML, GTR+F+R3, was determined using IQTree v2.1.4-beta^78^. Subsequently, the divergence times at the nodes of the phylogenetic tree using R8s v1.81, with calibration time points derived from the TimeTree database^79^. The calibration points included the divergence times between *M. truncatula* and *G. max* (55 million years ago, MYA), *M. truncatula* and *L. rhodostegia* (68 MYA), and *G. max* and *L. rhodostegia*(68 MYA). Finally, the CAFE v5.0^80^ program was employed to understand the expansion and contraction of gene families across the phylogenetic tree. The results indicate that *Z. insignis* (subfamily: Dialioideae) diverged from *S. tora* (subfamily: Caesalpinioideae) and *M. truncatula* (subfamily: Papilionoideae) around 51 MYA. The analysis of gene family dynamics identified 1,625 gene families with significant expansion (LRT test, *p*≤0.05) and 741 gene families with significant contraction (LRT test, *p*≤0.05) in the genome of *Z. insignis* (Figure 4A).

**Figure 4.**
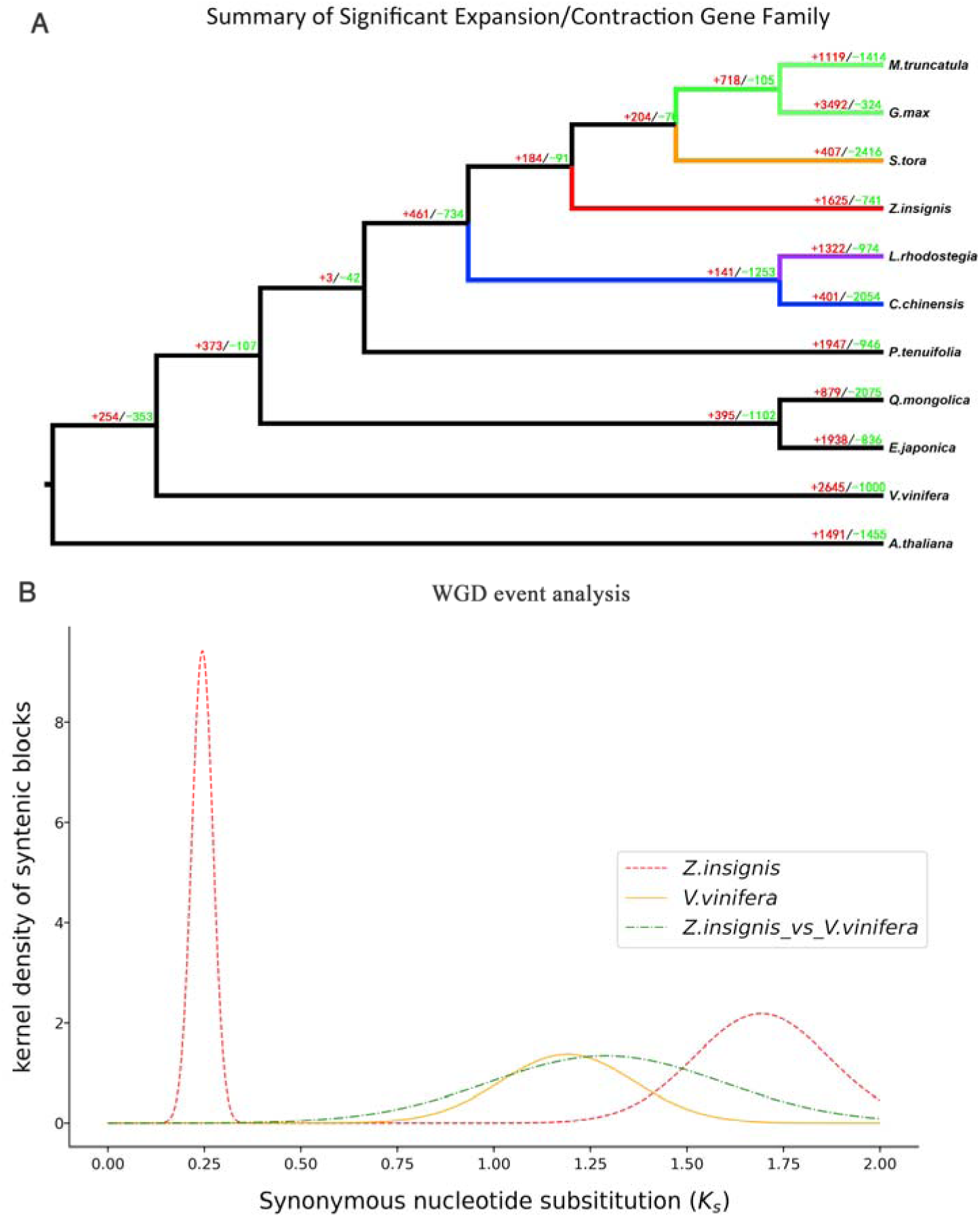
The result of gene family evolution and distribution of Ks values. (A) The result of gene family evolution. The plus/minus sign corresponding numbers reflect the expansion or contraction of gene families. The color of the branch indicates different subfamilies of Fabaceae. (B) The distribution of *Ks* values between *Z. insignis* and *V. Vinifera*. The *Ks* distribution curve of Z. insignis, V. Vinifera, and *Z. insignis* vs. *V. Vinifera* exhibited a significant overlap in the right *Ks* peak.

We conducted GO and KEGG enrichment analyses to understand the biological significance of the gene families that exhibited significant expansion and contraction with *a P-value* cutoff of 0.05. The GO and KEGG analyses were performed using the eggNOG-mapper web platform^59^. We employed TBtools v2.086^81^ to carry out the enrichment analysis of the identified gene families. The GO enrichment analysis revealed that the families that exhibited significant expansion were predominantly involved in molecular function and biological process (Supplementary Figure 2A). The terms of the molecular function are mainly ‘metal ion transmembrane transporter activity’, ‘inorganic cation transmembrane transporter activity’, and ‘cellulose synthase activity’ (Supplementary Figure 2A). And the terms of the biological process are mainly ‘hydrocarbon metabolic process’ and ‘olefin metabolic process’ (Supplementary Figure 2A). Furthermore, the KEGG enrichment analysis revealed that these families are significantly enriched in pathways such as ‘alpha-Linolenic acid metabolism’, ‘MAPK signaling pathway-plant’, ‘metabolism of terpenoids and polyketides’, ‘biosynthesis of other secondary metabolites’, and ‘ion channels’ (Supplementary Figure 2C). Conversely, for the families that exhibited a significant contraction, the GO enrichment analysis was predominantly involved in the biological process of ‘embryo development ending in seed dormancy’, ‘embryo development’, ‘seed development’, ‘fruit development’, and ‘proteolysis involved in protein catabolic process’ (Supplementary Figure 2B). The corresponding KEGG enrichment analysis identified pathways including ‘protein families: metabolism’, ‘ubiquitin mediated proteolysis’, ‘chromosome and associated proteins’, ‘protein processing in endoplasmic reticulum’, and ‘brite hierarchies’ (Supplementary Figure 2D).

### Identification of whole genome duplication events

To elucidate the whole genome duplication (WGD) events in *Z. insignis*, a comprehensive analysis was conducted using the WGDI toolkit v0.6.5^82^ with various parameters. We used *V. vinifera*^72^ as a reference genome. First, we used BLAST+ v2.15.0^83^ to identify homologous genes (e-value 10^-5^). We then drew a dot plot of collinear genes with the parameter -d (Figure 5A and B). Next, we conducted a collinear analysis with the parameter -icl, calculated *Ks* values with the parameter -ks, combined the results of the collinear analysis and *Ks* values with the parameter -bi, and fitted *Ks* peaks with the parameter -pf. Finally, we plotted a *Ks* dot plot with the parameter -bk (Figure 5C and D) and a *Ks* peak plot with the parameter -kf (Figure 4B). The dot-plot and *Ks* dot-plot results for *Z. insignis* showed a high degree of similarity to those shown in Figure 4A and C. They were cross-validated with the results from *V. vinifera* in Figure 5B and D. The *Ks* peak analysis revealed two peaks at 0.245±0.001 and 1.693±0.001, suggesting that *Z. insignis* likely experienced a lineage-specific WGD event, as illustrated in Figure 4B.

**Figure 5.**
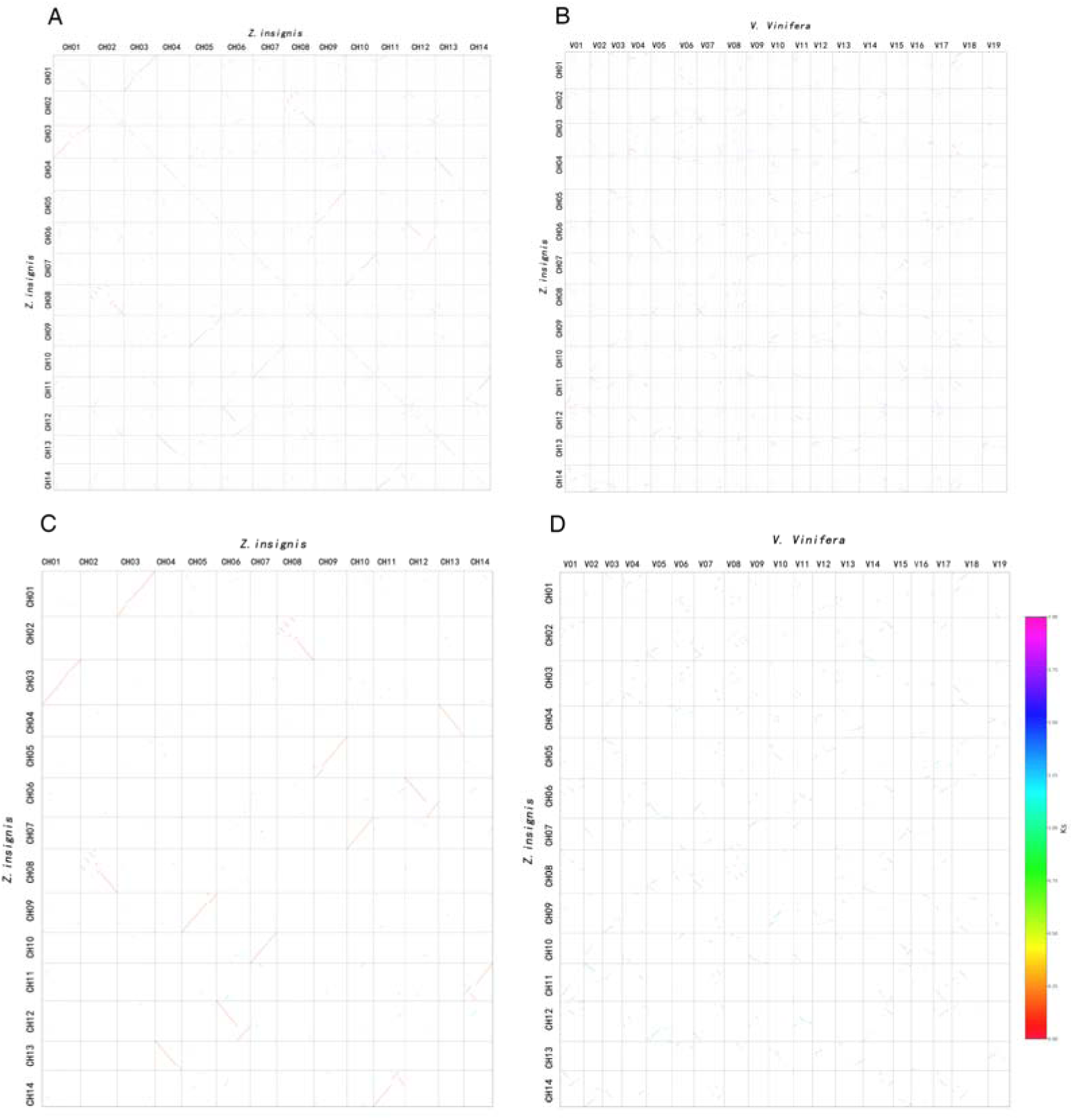
The result of WGDI analysis for *Z. insignis*. (A) The homologous genes dot plot of genomic synteny between chromosomes of *Z. insignis*. The best-hit genes are indicated in red, the second best-hit genes are in blue, and the other hit genes are in gray. (B) Dot plot of genomic synteny between *Z. insignis* and *V. Vinifera*. The best-hit genes are indicated in red, the second best-hit genes in blue, and the other hit genes are in gray (C) *Ks* dot plot of genomic synteny between chromosomes of *Z. insignis*. The color of the dot indicates the Ks of the gene pair. (B) Ks dot plot of genomic synteny between *Z. insignis* and *V. Vinifera*. The color of the dot indicates the *Ks* of the gene pair.

## Technical Validation

As part of our sequencing pipeline, we conducted a rigorous sample quality assessment for both DNA and RNA samples used in the study. The specific procedures and criteria for this assessment are detailed in our paper’s ‘Methods’ section, ensuring that only high-quality samples were selected for subsequent sequencing and analysis.

In our study, we employed two distinct methodologies to estimate the genome size of *Z. insignis*. The first method involved Flow cytometry analysis, which provided a quick and accurate measurement of the relative DNA content within the cells of *Z. insignis*. The second method was *K-mer* analysis, which leverages the frequency distribution of DNA sequences to predict genome size. The details have been mentioned in the ‘Method’ section. Both approaches were crucial in validating the genome size and ensuring the reliability of our genomic data.

In our assessment of the *de novo* genome assembly for *Z. insignis*, we utilized a suite of metrics, including N50, BUSCO, and LAI, to evaluate the performance of various assembly software tools. The results of these evaluations are presented in Tables 2, 3, and 4, which compare the assembly quality, completeness, and continuity across different assemblers. Based on these assessments, Falcon-purged output demonstrated the most favorable balance of completeness and continuity, making it the choice for subsequent downstream analyses. In summary, we have successfully generated a high-quality reference genome for *Z. insignis*, which may promote further biological and comparative genomic studies of this species.

## Code Availability

We used all software or pipelines, depending on the tools’ manuals or protocols. The software’s version is described in the methods section, and the parameters are given if not default.

## Acknowledgements

This research was supported by the Major Program for Basic Research Project of Yunnan Province (202401BC070001), the Youth Talents Special Project of Yunnan Province ‘Xingdian Talents Support Program’ (XDYC-QNRC-2022-0423), the National Natural Science Foundation of China, key international (regional) cooperative research project (No. 31720103903), the Yunnan Provincial Basic Research Program Youth Project (2019FD058), and the National Natural Science Foundation of China (No. 32270247). We are grateful to the Germplasm Bank of Wild Species, particularly the Molecular Biology Experiment Center and the iFlora High Performance Computing Center, the Kunming Botanical Garden, Ms. Ding-Jie Wang, and Mr. Zuo-Ying Xiahou.

## Author contributions

T.-S. Y. and H. L. Designed the experiments. S.-Y. C. and Z.-Y. Y. performed the experiments. S.-Y. C., H. L., and R. Z. analyzed the data. S.-Y. C., T.-S. Y. and H. L. wrote the paper. All Authors read and approved the manuscript.

---

**Supplementary Figure 1.**
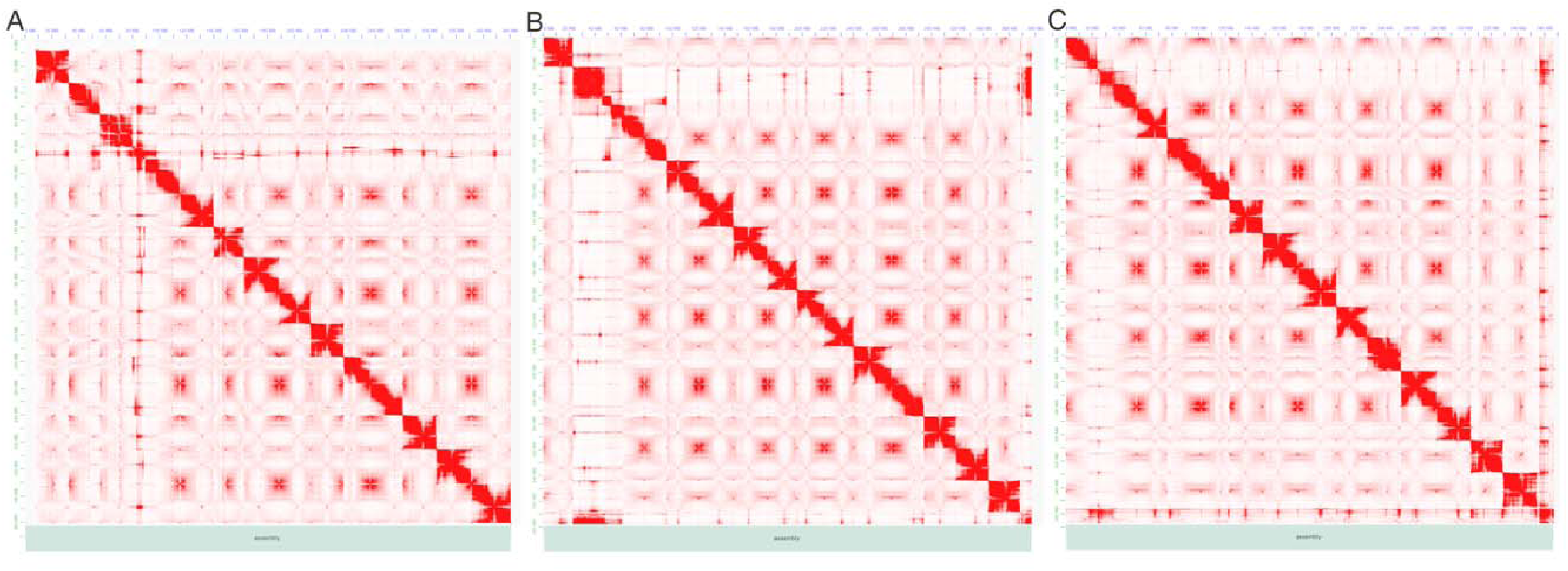
The heat map of the Hi-C interaction matrix generated by Purge_Dups. (A) Heat map of Hi-C for Falcon_purge. (B) Heat map of Hi-C for Canu_purge. (C) The Heat map of Hi-C for Flye_purge.

**Supplementary Figure 2.**
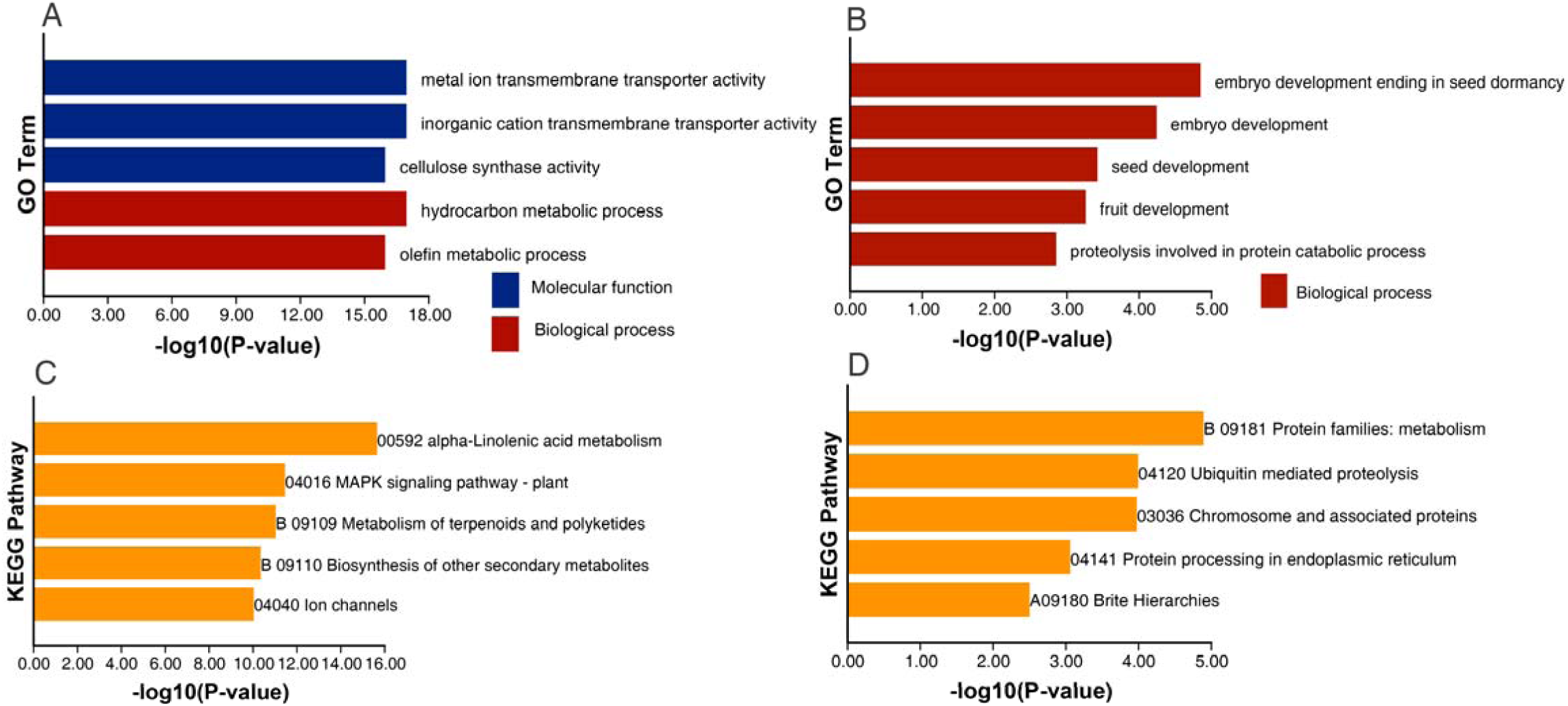
The result of enrichment analyses for GO and KEGG. (A)The result of GO enrichment analyses for significant expansion gene families (B) The result of GO enrichment analyses for significant contraction gene families. (C) The result of KEGG enrichment analyses for significant expansion gene families. (B) The result of KEGG enrichment analyses for significant contraction gene families.

